# Genome-wide Imputation Using the Practical Haplotype Graph in the Heterozygous Crop Cassava

**DOI:** 10.1101/2021.05.12.443913

**Authors:** Evan M Long, Peter J. Bradbury, M. Cinta Romay, Edward S. Buckler, Kelly R Robbins

## Abstract

Genomic applications such as genomic selection and genome-wide association have become increasingly common since the advent of genome sequencing. Genotype imputation makes it possible to infer whole genome information from limited input data, making large sampling for genomic applications more feasible, especially in non-model species where resources are less abundant. Imputation becomes increasingly difficult in heterozygous species where haplotypes must be phased. The Practical Haplotype Graph is a recently developed tool that can accurately impute genotypes, using a reference panel of haplotypes. The Practical Haplotype Graph is a haplotype database that implements a trellis graph to predict haplotypes using minimal input data. Genotyping information is aligned to the database and missing haplotypes are predicted from the most likely path through the graph. We showcase the ability of the Practical Haplotype Graph to impute genomic information in the highly heterozygous crop cassava (*Manihot esculenta*). Accurately phased haplotypes were sampled from runs of homozygosity across a diverse panel of individuals to populate the graph, which proved more accurate than relying on computational phasing methods. At 1X input sequence coverage, the Practical Haplotype Graph achieves a high concordance between predicted and true genotypes (R=0.84), as compared to the standard imputation tool Beagle (R=0.69). This improved accuracy was especially visible in the prediction of rare and heterozygous alleles. We validate the Practical Haplotype Graph as an accurate imputation tool in the heterozygous crop cassava, showing its potential for application in heterozygous species.

## INTRODUCTION

The past decade has seen an abundance of genomic sequence data produced for research and application in agricultural crops. With these new technologies, comes questions on how to effectively implement them (Torkamaneh *et al*. 2018). Two of the most common uses of genome-wide sequence data are genomic selection (GS) and genome-wide association studies (GWAS). While most GWAS attempt to locate distinct, causative regions of the genome, GS incorporates all available markers to predict traits (Meuwissen *et al*. 2001). Genomic selection leverages a training set population that has both genotypic and phenotypic data to predict traits in a related germplasm with only genotypic data (Heffner *et al*. 2009). This allows breeders to both increase accuracy when selecting traits with low heritability and to accelerate the rate of selections by decreasing cycle time (Xu *et al*. 2020).

While sequencing data has become increasingly common in agricultural applications, the financial cost remains a challenge to widespread implementation. Reduced representation marker systems have been produced to limit costs of performing genomic analyses (Romay 2018), all of which vary in marker density and depth, cost, and genotype confidence. In scenarios with limited diversity, such as single breeding pools or post-bottleneck populations, individuals share large stretches of sequence. The strong association between alleles in these blocks, or their linkage disequilibrium (LD), determines the number and distribution of genotype markers needed to explain the genetic variation in the population. High density of markers becomes more important when performing analyses in populations where LD decays quickly as in species with high diversity or among unrelated individuals. High marker density can also be beneficial to incorporate knowledge on previously studied loci across the genome.

To affordably obtain high density genotypes or to bridge information between different marker platforms it becomes necessary to impute missing genotypes from available genotype data. Increasing the stability across genotyping platforms and reducing per-sample costs becomes even more relevant in plant breeding scenarios, where many thousands of offspring are evaluated and changes in marker platform are common. Computational techniques to impute genome-wide information have been produced to bridge genotypic information from different marker panels and augment genotypic information from limited inputs (Yun *et al*. 2009). Genomic imputation methods often rely on a related training set with high confidence genotypic information to predict missing genotypes. These methods have been shown to improve consistency and efficiency of analyses of both genome wide associations (Spencer *et al*. 2009) and genomic selection (Cleveland *et al*. 2011).

Imputation is very common in genomic studies but is still plagued by barriers to high accuracy in many species. Known limitations of imputation stem from LD, allele frequencies, and population structure of the training population (Alipour *et al*. 2019). These difficulties are further compounded when working with a highly heterozygous crop, where both copies of the genome need to be modeled (Fragoso *et al*. 2016; Nazzicari *et al*. 2016). Heterozygosity introduces the challenge of phasing, or identifying which allele belongs to which copy of the genome, a challenge that is not limited to plants (Friedenberg and Meurs 2016). Imputation accuracy has been shown to affect the accuracy of genomic prediction in multiple scenarios (Pimentel *et al*. 2015; Wang *et al*. 2016; Van Den Berg *et al*. 2017). Highly accurate and less expensive imputation methods are needed to increase the gains made by GS by making genotyping more accurate and consistent. These improvements will enable research and breeding efforts to make accelerated gains, leading to more productive and adaptable crops in the changing global climate.

Rare variants contribute to the genetic load and overall performance of crops (Yang *et al*. 2017; Kremling *et al*. 2018; Kono *et al*. 2019), making high imputation accuracy, especially for alleles at low frequency, desirable for plant genomics applications. Diverse imputation tools exist and are often designed for different scenarios. One of the more common tools Beagle (Browning *et al*. 2018), which was designed for application in humans, works by leveraging LD between variants to predict missing genotypes. Beagle uses LD clustering to create an acyclic graph and a Hidden Markov model (HMM) to infer the most likely haplotype. Another method, EAGLE, leverages stretches of identity by descent (IBD) to perform long range phasing (Loh *et al*. 2016). In humans, where these imputation algorithms have been showcased, they have the advantage of large datasets with data from several thousands of individuals (Loh *et al*. 2016; Browning *et al*. 2018); this is not often possible in many plant breeding scenarios.

In maize, Beagle has difficulty accurately imputing rare variants, while a haplotype library based methods such as FILLIN can do so more easily (Swarts *et al*. 2015). A recently developed method known as the Practical Haplotype Graph (PHG) was created to leverage known haplotypes in a graph structure to efficiently impute genotypes (Bradbury, In prep). The PHG simplifies the genome to a set of distinct regions of the genome, for which it defines haplotypes. These haplotypes are constructed from whole genome sequence data or genome assemblies and are used to construct a trellis graph, capturing the diversity of haplotypes at each range and the relationships between adjacent haplotype regions. Sequence reads are then aligned to the graph and a HMM is applied to predict the most likely haplotypes. By aligning reads to pan-genome haplotypes, the PHG minimizes errors due to reference bias, poor alignment, and mis-called variants.

Here we showcase the potential application of the PHG in imputation of heterozygous crops. The PHG has already been shown to be an efficient tool for aiding imputation and genomic selection in breeding of the inbred cereal crop Sorghum (Jensen *et al*. 2020). It has also been implemented to impute genotypes in highly diverse maize lines (Franco *et al*. 2020). To show the utility of the PHG in a heterozygous crop we must overcome two distinct challenges: (1) obtaining phased haplotypes to populate the database and (2) modeling both copies of the genome accurately. Without an abundance of data, it is very difficult to obtain accurate phasing in a highly heterozygous species. This study will explore these challenges by imputing haplotypes from low-coverage skim sequencing, while comparing results to Beagle’s performance.

To investigate the construction and performance of the PHG in a heterozygous scenario, we created a PHG for cassava (*Manihot esculenta*), a root crop with high levels of heterozygosity reinforced by centuries of clonal propagation. Cassava is a major caloric source for over half a billion people around the world, with a high concentration in sub-Saharan Africa (Parmar *et al*. 2017). Improved imputation in cassava could enable greater gains in breeding efforts to increase food security. In this study we utilize sequence data from the previously published HapMapII in cassava (Ramu *et al*. 2017), which includes whole genome sequence (WGS) data for 241 cassava clones. This data is used to produce a PHG in cassava and illustrate its effectiveness in genomic imputation in a heterozygous crop. We further validate these methods through genomic prediction and simulation.

## MATERIALS AND METHODS

### Haplotype Sampling

Genomic data was used from the second-generation Cassava Haplotype map consisting of 241 taxa, including both cultivated and wild germplasm (Ramu *et al*. 2017). Raw data is composed of short-read, whole genome sequence data from each taxon amounting to greater than 20X coverage on average. The high depth of the sequence data is necessary to accurately distinguish between heterozygous and homozygous variants. Haplotype regions, termed here as reference ranges, were defined by genic regions with additional 500bp flanking sequence from the Cassava V6 reference genome.

The detailed process of creating a PHG is outlined at “https://bitbucket.org/bucklerlab/practicalhaplotypegraph/wiki/Home” and has been described previously (Jensen *et al*. 2020; Franco *et al*. 2020). Here, we outline the specific steps taken to create a PHG in the heterozygous crop cassava. The major hurdle to producing a haplotype graph in a heterozygous species is obtaining accurately phased haplotypes. Because many of these cassava lines are cultivated taxa, we expected to find identical by descent (IBD) haplotypes brought about by generations of breeding within restricted breeding pools. These IBD segments provide confidently phased haplotypes as well as capturing their relationships to adjacent haplotypes (Fig. 1). We identified and sampled these homozygous haplotypes which we inferred to represent IBD haplotypes. This was done by measuring the number of heterozygous variants for each reference range in each taxon, then classifying those haplotypes as homozygous or not. The threshold for haplotypes to be considered IBD was determined empirically to be 0.001 heterozygous SNPs per base pair (Supplemental Fig. 1), as *de novo* mutations or errors in variant calling may produce low levels of perceived heterozygosity. This threshold was additionally validated by testing imputation accuracy of the PHG.

**Figure 1.**
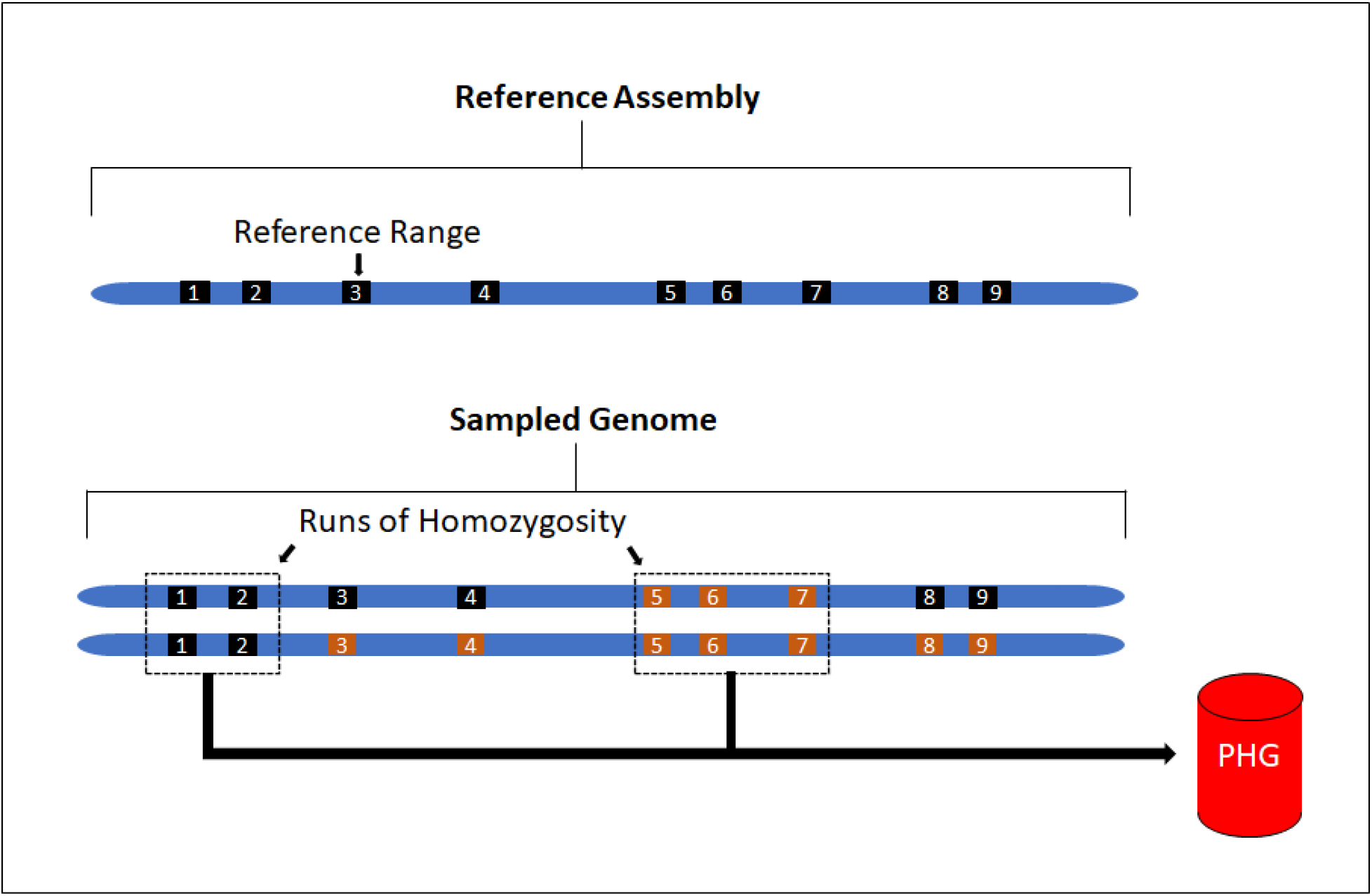
Haplotype view of the genome. Top) Representation of reference ranges informed from genic regions from the reference genome. Bottom) Haplotypes sampled from runs of homozygosity for use in PHG with different colors representing separate haplotypes at a given region (i.e. ranges 1,2,5,6,7 are homozygous and haplotypes can be sampled).

After haplotypes were sampled from IBD regions of the genome, they were loaded as GVCF files into a PHG database. Similar haplotypes were then collapsed based on sequence similarity to produce a representative set of available haplotypes. Haplotypes are collapsed to make alignment more efficient, while retaining as much distinct haplotype information as possible. Collapsing is performed using an unweighted pair group method with arithmetic mean (upgma) tree from pairwise distance matrix from sequence variants to measure the similarity between haplotypes. Based on imputation accuracy tests, we chose a level of similarity (PHG parameter: maximum divergence) to collapse haplotypes of 0.001, corresponding to less than 1 in 1000 nucleotide differences between haplotypes. This level of collapsing maintains high accuracy while collapsing redundant haplotypes (Supplemental Fig. 2). We then produced a pan-genome composed of consensus haplotypes representing the diversity of haplotypes.

### Predicting Haplotypes

Once we obtained a set of consensus haplotypes, we implemented an HMM to infer genome-wide haplotypes from low depth genotyping data. Sparse genotype information was created by downsampling whole genome sequence data randomly using samtools to simulate skim sequencing. We randomly sampled 20 taxa from the cultivated varieties within the population to serve as a test set for downstream analyses. To test different levels of sequencing depth, we down-sampled reads to amounts estimated to represent 0.1X, 0.5X, 1X, 5X, and 10X single-end, whole genome sequence coverage.

These sampled sequences were aligned to the consensus haplotypes stored in the PHG to impute whole genome variants. A trellis graph is formed with every reference range representing separate ranges and the consensus haplotypes as nodes at each of those ranges. The most likely paths through the graph were then determined using an HMM Viterbi algorithm. Because cassava is heterozygous and diploid, this step produces the two most likely paths for each taxon. The emission and transition probability parameters of the HMM are defined by the genomes of the reference population used to build the database. The emission probabilities are calculated by considering the probability of two given haplotypes, given the aligned reads. The transition probabilities are defined by the edges between haplotypes in the PHG.

Due to the sparse sampling of IBD haplotypes from heterozygous taxa used to produce the PHG, the database lacked abundant transition information between adjacent reference ranges. To compensate for this, we aligned WGS for all 241 taxa used to create the database and predicted most likely paths through the graph. These paths were then used to augment the transition probabilities, without contributing any additional haplotypes.

### Beagle imputation

We compared our imputation accuracy results to the common genotype imputation tool Beagle (Browning *et al*. 2018). Beagle was developed for the purpose of human data, but is a common tool used by many plant studies to impute missing genotypes. Because Beagle v4 can incorporate genotype likelihoods based on read depth, we used it for the imputation of the low depth sequence when it improved accuracy, otherwise we utilized Beagle v5. We used the same HapMapII data from the 241 clones to impute missing genotypes with Beagle.

### Genomic Prediction

We used 57 clones from a single breeding program, to reduce effects of population structure, to determine the impact of imputation errors on genomic prediction accuracy using cross validation. Reads were down-sampled and imputed as previously described. Three root traits were used for genomic cross validation: fresh root yield, root size, and root number. Phenotypes for each clone were downloaded from CassavaBase.org, constituting 57 clones, spanning 23 years from 1996 to 2018, across 13 locations in Africa. Ten-fold cross validation was performed by randomly selecting 10% of the clones to hold out and predict using the remaining clones as a training set. The correlation between predicted phenotype and the observed BLUE was used as the prediction accuracy. We performed 50 replications as well as a single holdout prediction to measure genomic prediction accuracy. A single step model was performed:

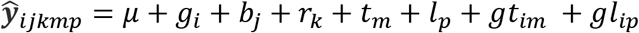

Here, 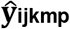 is the predicted trait and **μ** is the fixed effect of the overall mean. Random effects were fitted as follows: **g_i_** is genotype effect of the i^th^ clone, **b_j_** is the effect of the j^th^ block, **r_k_** is the effect of the k^th^ replicate, **l_p_** is the location of the p^th^ location, **t_m_** is the effect of the m^th^ year, **gl_ip_** is the interactive effect of the i^th^ clone and the p^th^ location, and **gt_im_** is the interaction effect of the i^th^ clone and the m^th^ year. This was performed using the mixed model tool Echidna (Gilmour 2019).

The vectors of random effects for the model were distributed as

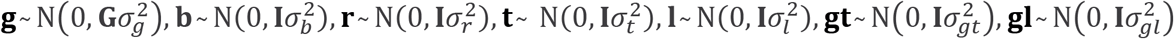

Where **G** is the genomic relationship matrix calculated using the “Eigenstrat” method of the R package SNPRelate (Zheng *et al*. 2012) and **I** is the identity matrix.

### Pre-phased Haplotype PHG

We investigated the viability of using computationally phased haplotypes to curate a PHG database rather than relying on IBD regions of the genome. First, we phased the variants from the 241 cassava clones using a combination of Beagle (Browning *et al*. 2018) and HAPCUT2 (Edge *et al*. 2017). These variants were used to create a PHG to be tested against the IBD version of the PHG. The second test utilized Oxford Nanopore (ONP) long-read sequencing from six cassava clones within the HMII population. High molecular weight DNA was extracted from young cassava leaves, selected for fragments 20-80 Kb long, and sequenced with MinION following the manufacturer recommendations. Variants were called using Guppy and their variants phased with WhatsHap (Schrinner *et al*. 2020). These six clones were then used to populate another PHG, we will identify as the “ONP6 PHG”. Larger reference ranges were divided into smaller regions to increase the probability of sampling correctly phased haplotypes. Twenty clones with the highest relationship to the six taxa with ONP data were used as the test set for these tests.

### Imputation from Simulated Genotypes

A sample of 20 related individuals from the HapMapII population were selected to serve as parents for a simulated genotyping scenario. The genomes were phased using Beagle and then used to populate a PHG database. We then used these parents to simulate 5 generations of random mating given a population size of 100 (Supplemental Figure 3.). Forward genetic simulations were completed using SliM (Haller and Messer 2019). Artificial short read-sequencing was then simulated for these offspring using neat-genreads (Stephens *et al*. 2016) at varied coverage levels. Reads were then aligned using bwa used to call and impute variants using Sentieon (Kendig *et al*. 2019) and Beagle. Reads were also aligned to the PHG formed from the original parents for imputation.

### Data Availability

Supplementary files and scripts used for the production and testing of the cassava PHG can be found at https://bitbucket.org/bucklerlab/p_cassava_phg. Genotype and phenotype data from HapMapII (Ramu *et al*. 2017) was downloaded from cassavabase.org. Support and methods for practical haplotype graph implementation can also be found at https://bitbucket.org/bucklerlab/practicalhaplotypegraph/wiki/Home. Raw Oxford nanopore sequence data for this project is available at NCBI BioProject ID PRJNA589272.

## RESULTS

### Haplotype Sampling

To obtain phased haplotypes for the PHG we sampled haplotypes from homozygous regions of each clone. Centuries of clonal propagation and reported inbreeding depression (de Freitas *et al*. 2016) suggest cassava germplasm would be highly heterozygous, however, we found that, on average, ~20% of all reference ranges from each taxon were homozygous. This resulted in a high number of missing haplotypes in each taxon, but a high confidence in the phased haplotypes that were sampled. Despite the variability in the number of homozygous samples by reference range, >90% of the reference ranges were homozygous in at least 10% of the HapMapII population (Supplemental Fig. 4). From these IBD haplotypes we were able to sample ~50% of the segregating sites. This proportion increased to 77% when considering sites with minor allele frequency above 5%, suggesting that many of the common variable sites have been sampled.

### Imputation and Genomic Prediction Accuracy

Because imputation accuracy is dependent on the relative allele frequency and phase of the allele being called, we classified genotype calls by allele frequency class: homozygous major, homozygous minor, and heterozygous. In our analyses, imputation accuracy is defined as the ability of the imputation method to reconstitute genome-wide SNPs from the input data. We use the correlation between the predicted alleles and the true alleles (defined by HapMapII) as a metric to make the PHG and Beagle comparable, because the PHG utilizes reads and Beagle utilizes variants to make their predictions.

Imputation of skim sequence genotyping showed PHG methods had a large advantage over Beagle using low coverage sequence. At a level of 1X coverage random sequencing, the PHG predicted allele calls with a correlation of R^2^=0.84, while the correlation between Beagle predicted alleles and the true calls was R^2^=0.69 (Fig. 2 A). At higher depths of coverage (>5X), the raw data provides ample information to distinguish between homozygous and heterozygous genotypes, allowing Beagle to determine the correct genotype. The PHG, however, is able to distinguish between the available haplotypes at a coverage of 0.5X and adding additional sequence data does not increase the accuracy, as there is no correlation between accuracy and coverage beyond 0.5X.

**Figure 2.**
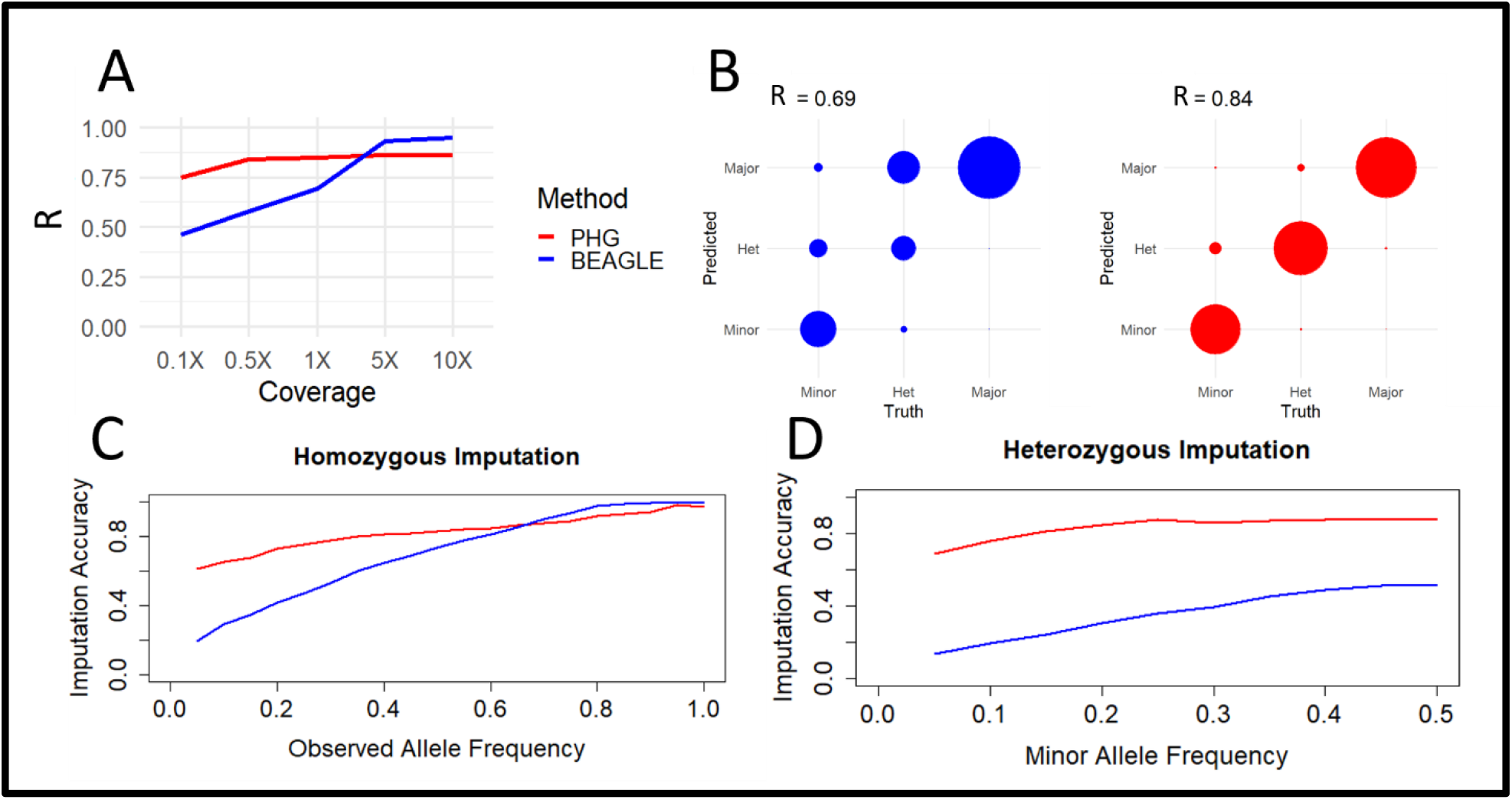
Imputation accuracy from skim sequencing. A) Displays correlation between imputed and true variants by imputing with the PHG and Beagle at different levels of skim sequencing. B) Displays concordance between true and imputed allele at 1X coverage separated by alleles classes: minor, heterozygous, and major. C) Imputation accuracy at 1X coverage is shown for homozygous genotypes separated by allele frequency of the true allele a that locus. D) Imputation accuracy at 1X coverage is shown for heterozygous genotypes separated by minor allele frequency at that locus.

The improved performance of the PHG is most noticeable in its accurate predictions of heterozygous and rare genotypes. The PHG was able to impute genotypes with high accuracy regardless of allele class (Figure 2B). The PHG’s high accuracy at low allele frequencies for both homozygous (Figure 2C) and heterozygous genotypes (Figure 2D), display its ability to impute rare alleles.

The imputed genotypes were then utilized in a genomic prediction scheme consisting of 57 cassava clones (Supplemental Fig. 5) from a single breeding program. The related nature of the clones ensured an adequate level of heritability to assess genomic prediction accuracy. Ten-fold cross validations and single holdout validation showed that imputation accuracy generally appeared to follow the genomic prediction accuracy, for fresh root yield and root number, while no clear pattern was apparent for the root size trait (Fig. 3).

**Figure 3.**
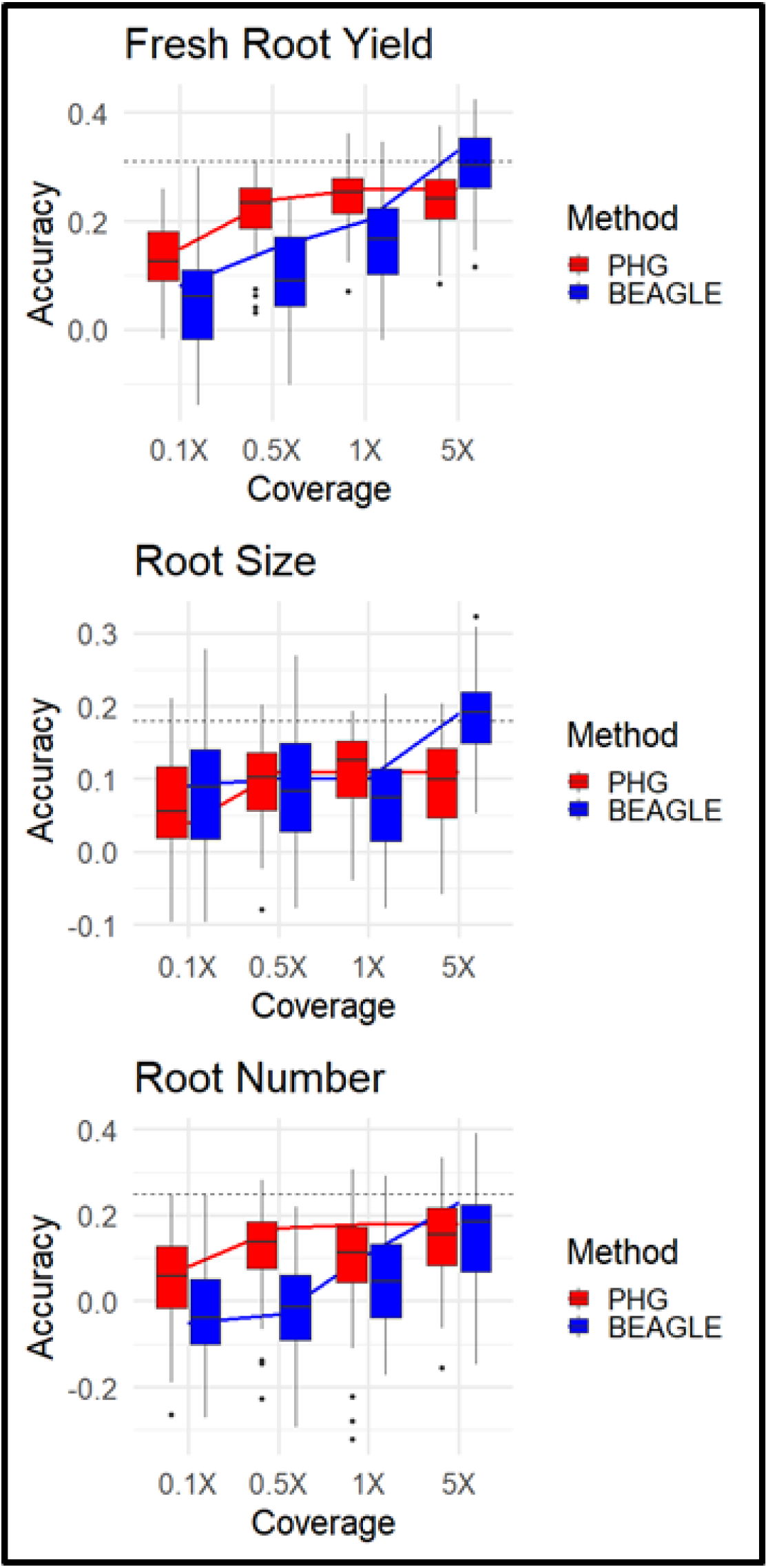
Genomic Prediction Cross Validation. 10-Fold cross validation (box) and single holdout cross validation (line) show genomic prediction accuracies of 3 root traits using different imputation methods at varied sequence depths. Single holdout cross validation using complete genotype dataset is shown (dashed line).

### Phased Haplotype PHG

We tested the viability of populating the PHG with haplotypes phased by other methods. We compared the IBD method of sampling phased haplotypes to two methods of phasing variants. The first method used Beagle and HAPCUT2 to phase the variants called from the HapMapII WGS data. The second method utilized six cassava clones with ONP long-read data. The IBD method of populating the cassava PHG produced the highest accuracy (Fig. 4). These results suggest that Beagle and HAPCUT could not accurately phase heterozygous haplotypes at this scale but maintained comparable accuracy due to IBD haplotypes. While the PHG was made from 6 clones with ONP data, it relied on a far narrower set of germplasm. This suggests that accurate haplotypes are likely captured using this method but lack adequate sampling to capture sufficient haplotypes.

**Figure 4.**
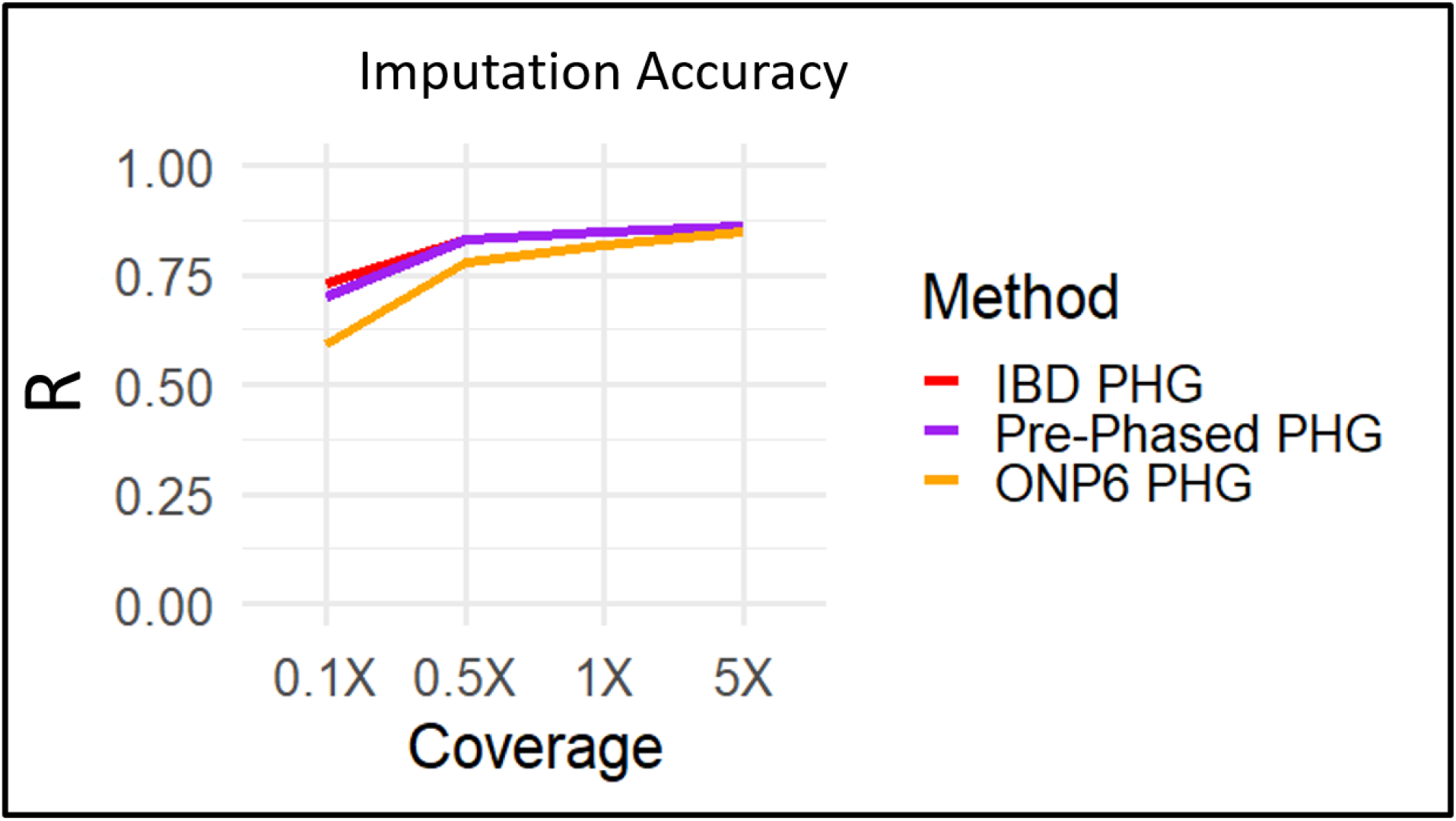
Haplotype phasing methods in the PHG. Imputation accuracy is shown for three different methods of populating a PHG. First the IBD PHG (red) was populated using homozygous haplotypes from the 241 HapMapII clones. Second, the Pre-Phased PHG (Purple) used Beagle and HPACUT2 to phase these same clones. Third, the ONP6 PHG (Yellow) used ONP long-reads and WhatsHap to phase six related taxa to the test set.

### Imputation Simulation

Evident from the tests using haplotypes from IBD regions of the genome, sampling phased haplotypes is a difficult aspect of creating an effective PHG in a heterozygous species. To explore the performance of the PHG in a scenario where one could aptly sample the diversity of haplotypes, we used simulated offspring from a set of 20 phased genomes. While phasing errors exist, we accepted these phases as truth for the simulation of offspring. This ensured that all haplotypes present in the offspring exist in the PHG database. We found that the disparity in accuracies between PHG and Beagle at high sequence coverage disappeared in our simulation (Fig. 5), while the trend in Beagle accuracy was very similar to our empirical tests. While the simulation does represent an ideal scenario, including a narrower set of germplasm, it highlights the performance of the PHG when accurately phased haplotypes are available.

**Figure 5.**
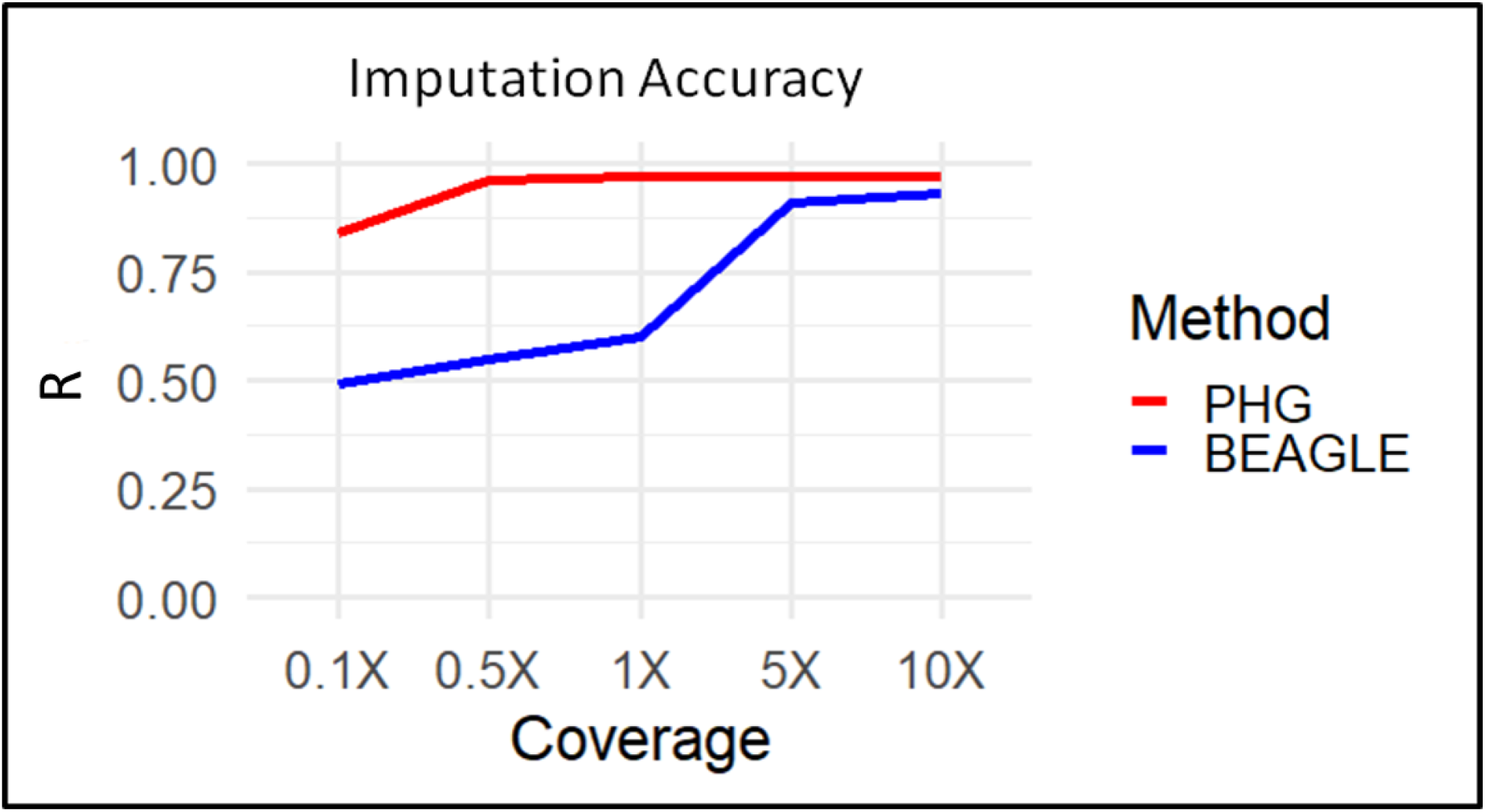
Imputation accuracy with simulated genotypes. A simulated scenario where a PHG is populated using 20 parents with full phased information. Correlation between imputed and true variants by imputing with the PHG and Beagle at different levels of skim sequencing.

## DISCUSSION

We have detailed a method of implementing a PHG for the heterozygous plant species cassava. This PHG database utilizes phased haplotypes to predict missing genotypes from low depth input sequence. Runs of homozygosity formed by IBD relationships proved to be the most reliable method of sampling phased haplotypes given the available data (Fig. 4). This method of obtaining haplotypes, while not able obtain the full diversity of alleles, captures 77% of common alleles and produces ample haplotypes for significant imputation accuracy at very low sequence depth (Fig. 2A).

The high accuracy of the PHG demonstrates its potential as an imputation tool for use in heterozygous crops. The advantages of the PHG imputation methodology are especially evident in its accuracy at calling rare and heterozygous alleles (Fig 2C,2D). Furthermore, the observed weaker relationship between allele frequency and imputation accuracy, highlights its ability to predict rare alleles. Across both simulated and empirical experiments, we found that the ability of the PHG to impute whole genome variants was consistent at or above 0.5X sequence coverage. The haplotype-based representation of the genome enables this imputation methodology to overcome the logistical hurdles such as those produced by sequencing and assembly errors, repetitive sequences, and poor alignments.

The plateau reached in imputation accuracy (Fig. 2A) using the PHG most likely indicates that we have not sufficiently sampled the diversity of possible haplotypes. At sequence coverages of 5X and more, the raw data can produce the true genotypes and little imputation of missing genotypes is occurring. The disparity between the PHG and Beagle at these high coverages shows the presence of missing haplotypes in the database, rather than any disparity in performance. This is supported by a visible relationship between homozygous incidence in our population and reference range imputation accuracy (Supplemental Fig. 6), suggesting that those ranges with poor imputation accuracy were not amply sampled. The length and abundance of the IBD runs of homozygosity in our dataset likely determine the ability of the HMM to accurately predict haplotypes. This hypothesis for decreased accuracy is also supported by the removal of such a disparity under simulation, where all possible haplotypes are sampled in the database (Fig. 5). These results suggest that, although an already powerful tool, the PHG achieves maximum performance with sufficient sampling of available haplotypes.

While the imputation accuracy of the PHG is limited based on the haplotype sampling, its high accuracy with low levels of input sequence highlights its potential for genomic applications, where sparse genotyping is common. We showed that this is true regarding genomic prediction by performing cross-validations with the imputed genotypes (Fig. 3). The genomic prediction was still limited by imputation accuracy, but by enabling higher accuracy we can achieve more reliable predictions (Pimentel *et al*. 2015; Wang *et al*. 2016; Van Den Berg *et al*. 2017).

With increased environmental pressures in a changing global climate and growing populations, accelerated breeding is vital to sustainable food production. With increased imputation accuracy from more limited genotyping inputs, smaller breeding programs can afford to implement GS, enabling them to increase selection pressure across their breeding pools. Similarly, imputation to genome-wide scale can bridge gaps between different data sets containing information on different marker panels, enabling the use of larger datasets for prediction. Accurate imputation could also enable breeders to utilize genomic prediction models that incorporate more prior functional information on genome-wide variant effects into predictions, using methods such as GFBLUP (Fang *et al*. 2017) or BayesR (MacLeod *et al*. 2016; Van Den Berg *et al*. 2017). These possible applications of imputation have the potential to increase total genetic gain made by breeding programs.

We show that while computational methods might not be able to solve haplotype phasing with short-read data, long-read sequencing may be able to overcome that issue. While limited in scope, the ability of the PHG created from six clones with ONP data suggests the potential application of long reads for obtaining phased haplotypes. One could envision a breeding scenario in which parents are sequenced and phased using long-reads and offspring are predicted from minimal genotyping input using the PHG. Then every few generations shallow WGS can be used to update the PHG and compensate for changing LD structures.

## CONCLUSION

The PHG is a method to reduce a genome to a set of haplotypes. We have shown that this method can predict whole genome haplotypes in a heterozygous species from sparse genotyping information. Its high accuracy, especially in rare alleles, at very low depths of skim sequencing makes it a potentially powerful imputation tool. Continued work in populating the PHG database with confidently phased haplotypes will lead to a more consistent prediction model across varied genotyping methods.

## ACKNOWLEDGMENTS

This study is made possible by the funding and support of the Nextgen Cassava project, the Bill and Malinda Gates foundation, and the USDA-ARS. We also acknowledge the programming staff in the Buckler lab who created and support the development of the practical haplotype graph. Lastly, we are grateful for the greater Nextgen Cassava community for supporting the curation of genotype and phenotype data used in this project as well as the organization of this data in cassavabase.org.

## SUPPLEMENTAL FIGURES

**Supplemental Figure 1.**
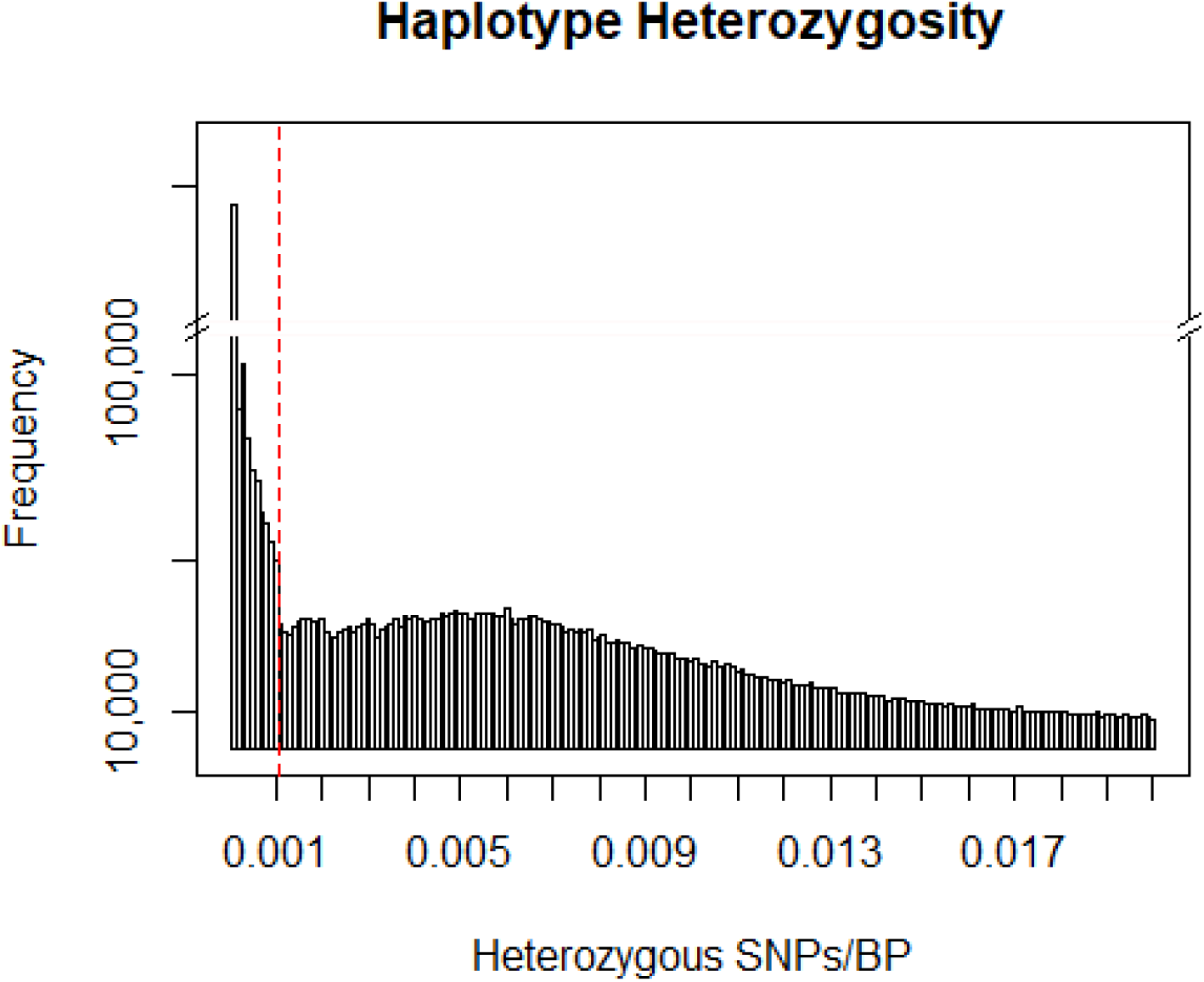
Histogram of sampled haplotypes by the number of heterozygous SNPs per base pair. Dotted line shows the threshold chosen to distinguish nearly IBD haplotypes.

**Supplemental Figure 2.**
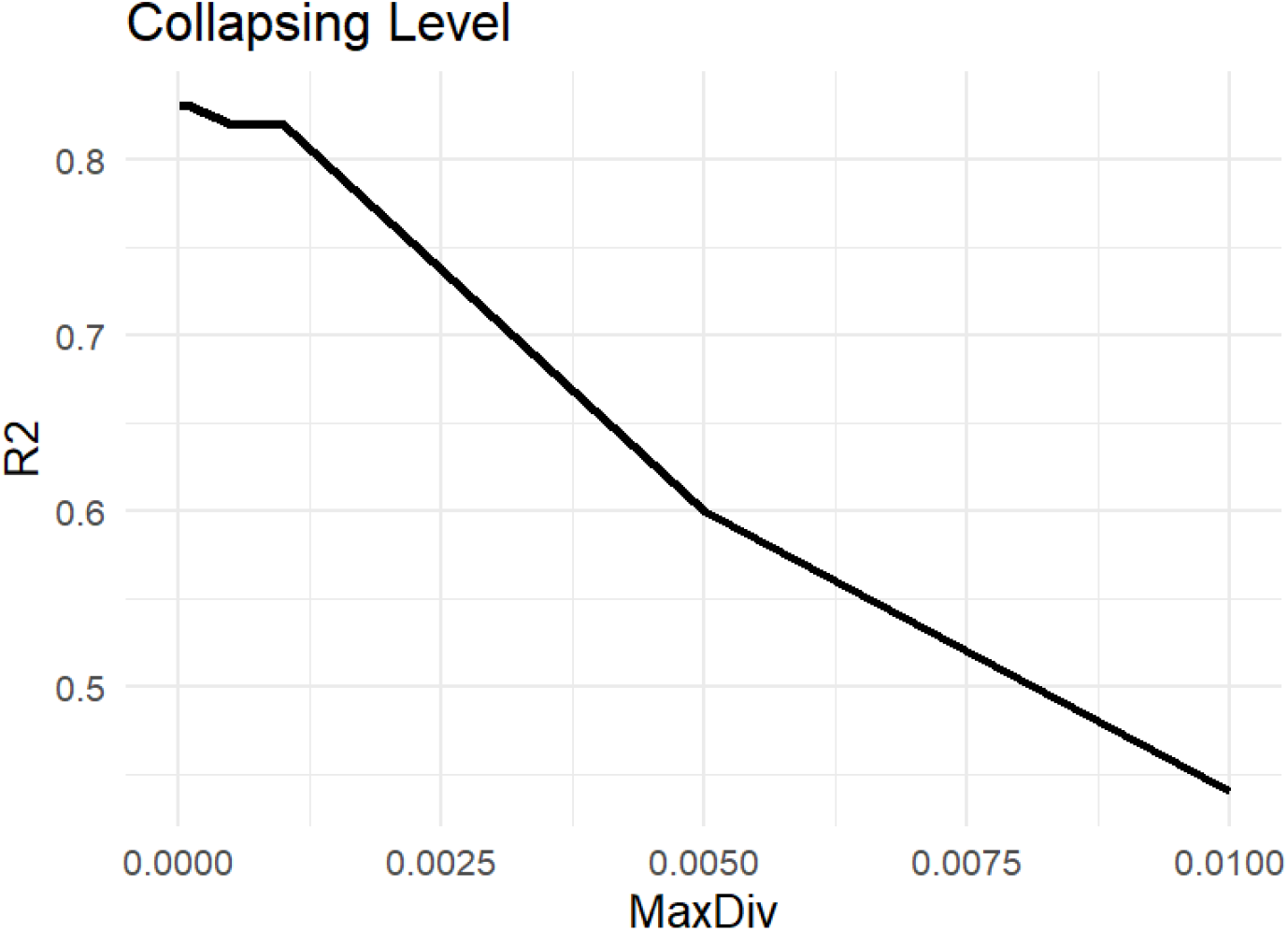
Correlations of Imputed calls to true calls at different haplotype collapsing levels based on the maximum divergence parameter of the PHG.

**Supplemental Figure 3.**
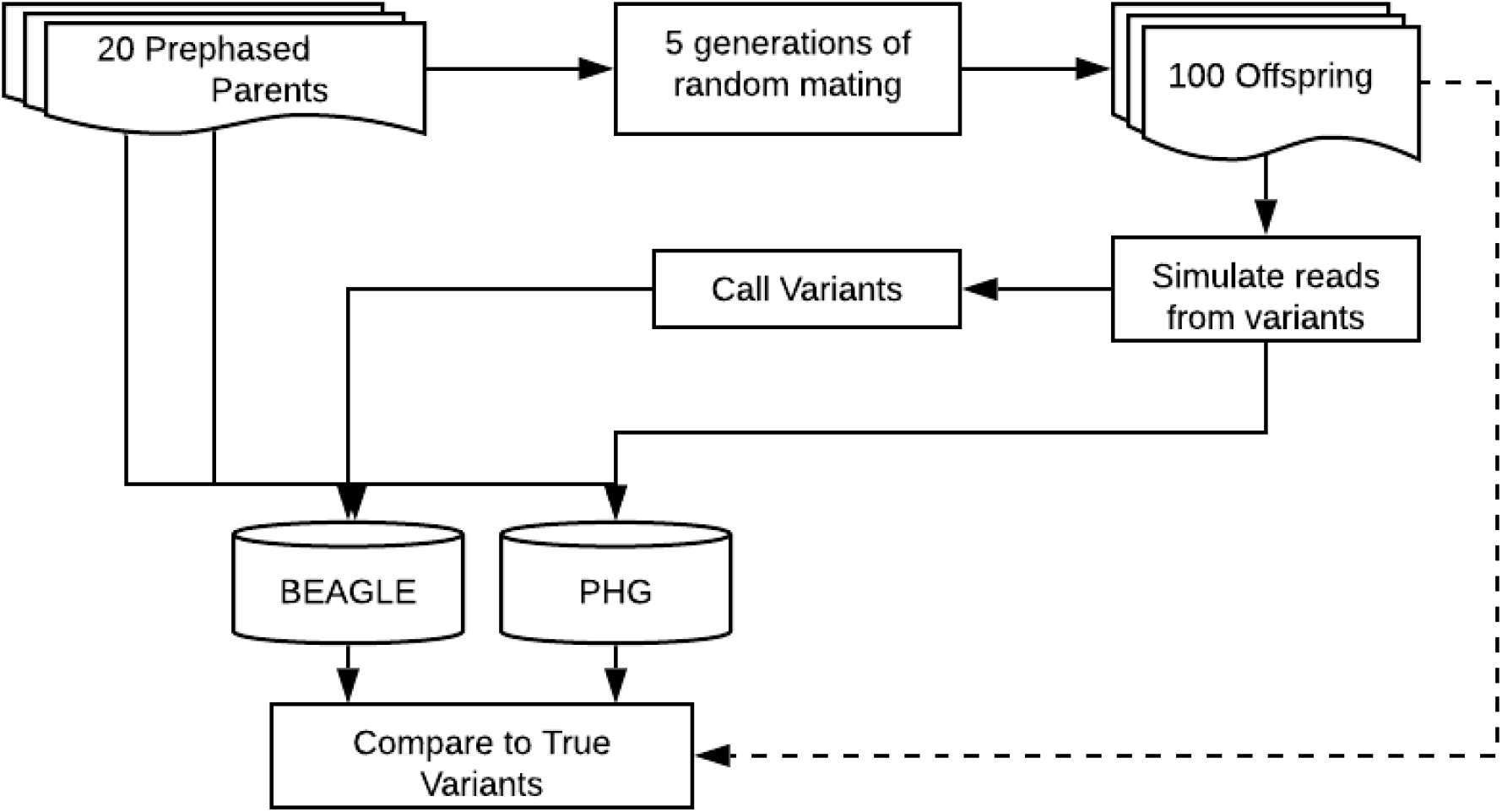
Simulation and Imputation Schema. Phased genomes from 20 parents from single breeding program were used to simulate generations of mating and recombination.

**Supplemental Figure 4.**
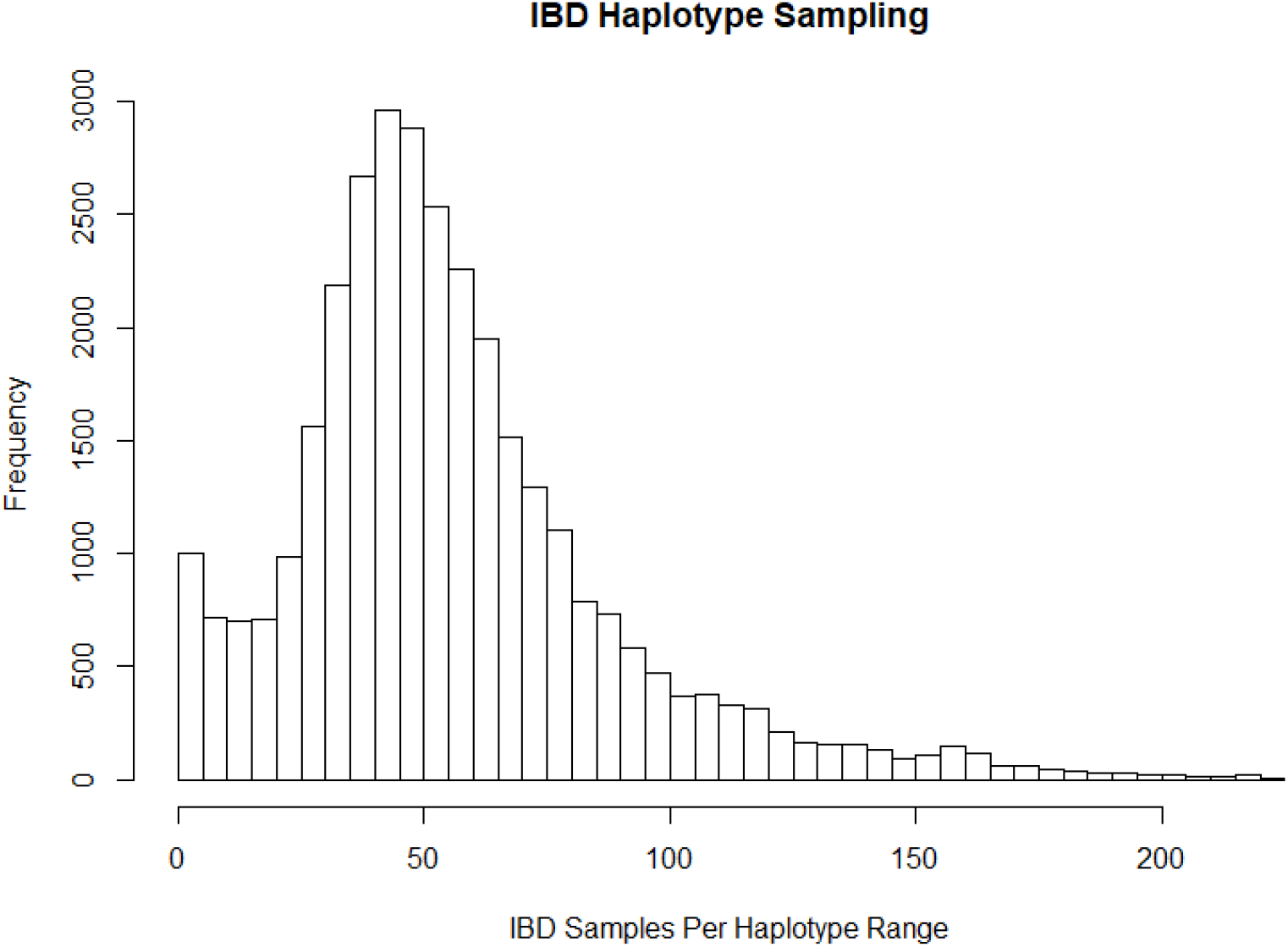
Histogram of IBD sampling frequency of all reference ranges. Y-axis shows the number of reference ranges with a given number of IBD samples.

**Supplemental Figure 5.**
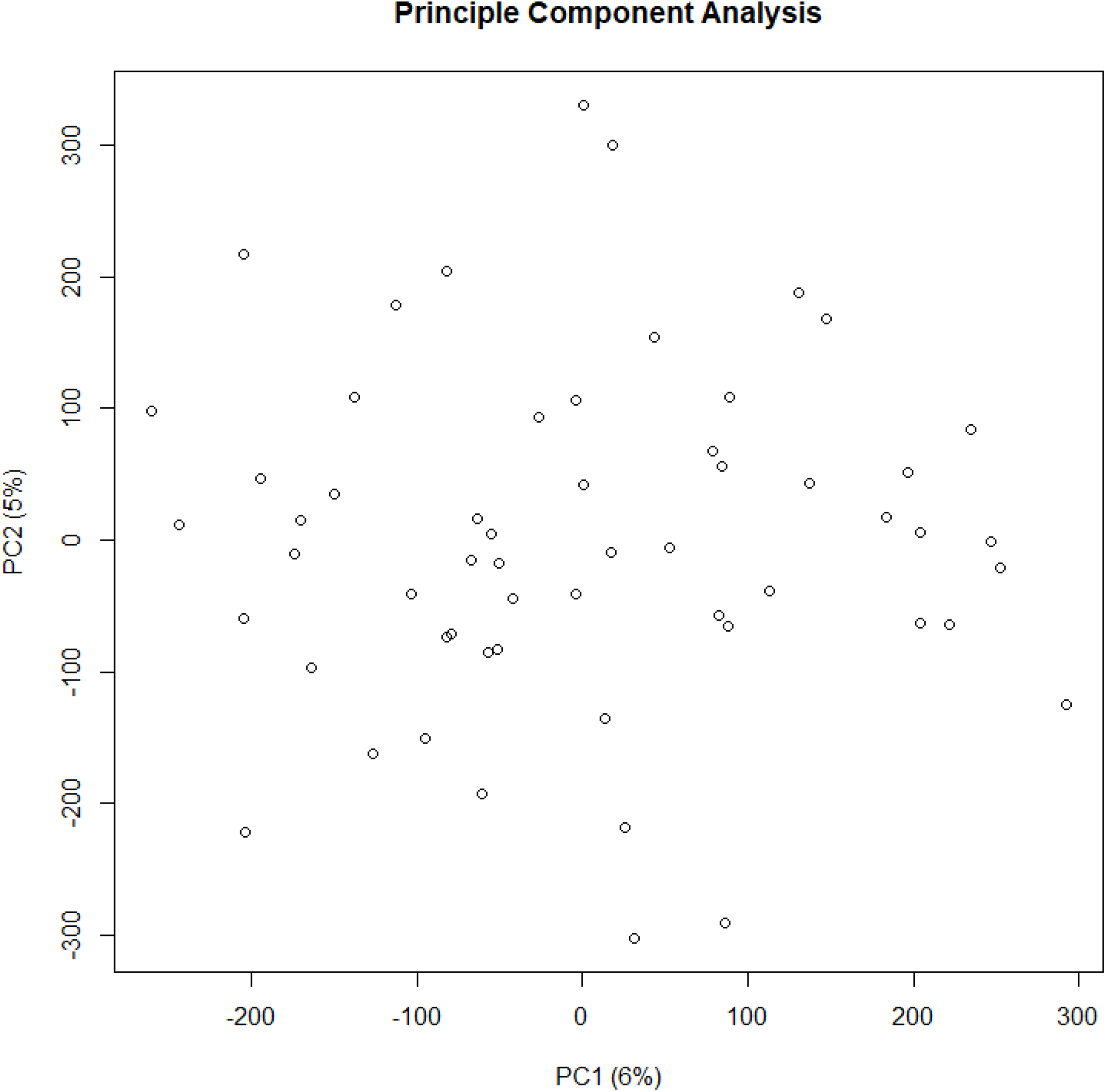
Principal component analysis of 57 clones used in genomic prediction cross validation. Lack of clusters show little population structure among the clones.

**Supplemental Figure 6.**
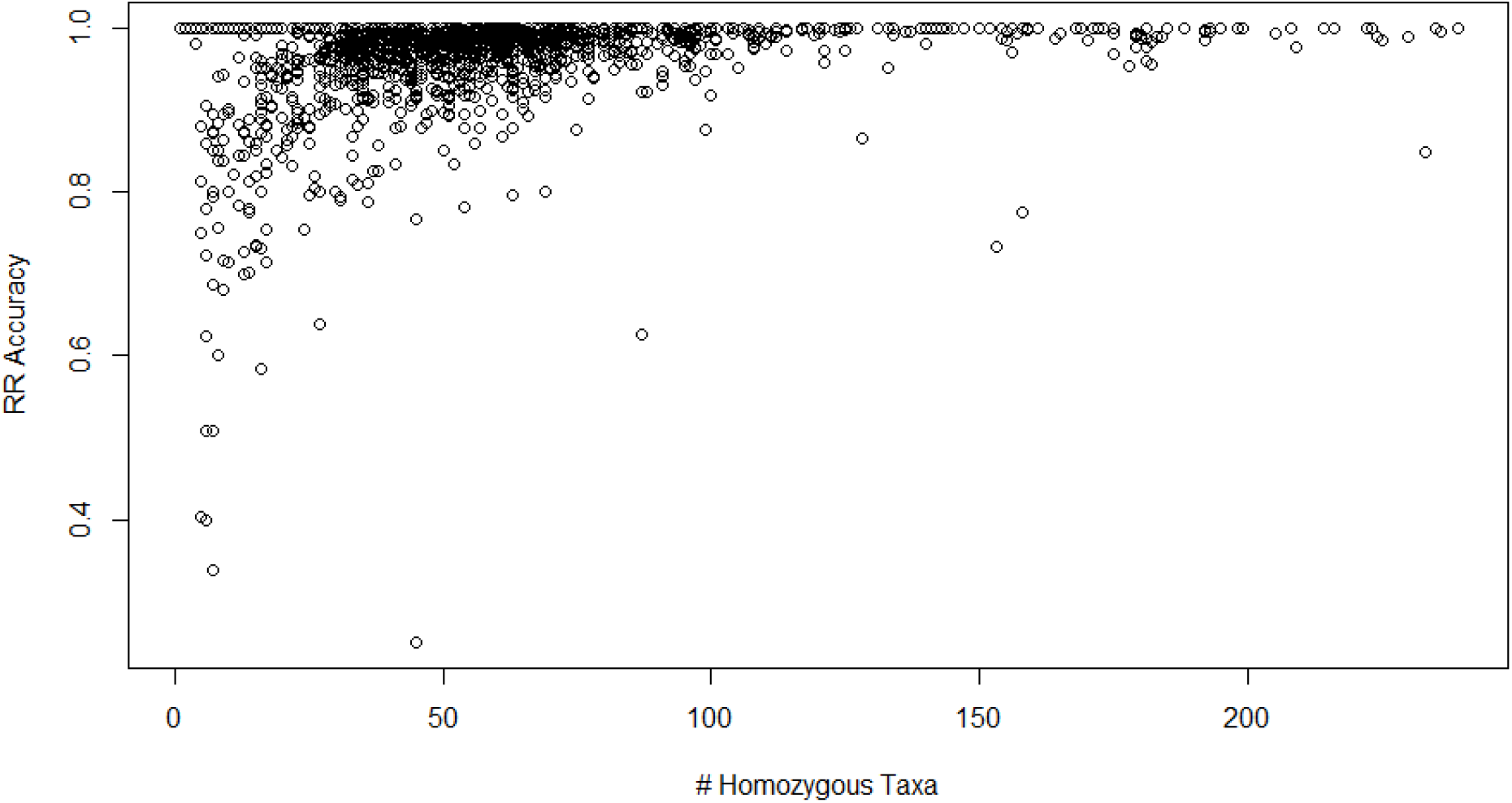
Imputation accuracy at each reference (y-axis) range by homozygous incidence in the HapMapII Population (x-axis). Low accuracy shown at reference ranges with low incidence of homozygous taxa.

